# Single cell transfection of human induced pluripotent stem cells using a droplet-based microfluidic system

**DOI:** 10.1101/2020.06.09.143214

**Authors:** C. Pérez, A. Sanluis-Verdes, A. Waisman, A. Lombardi, G. Rosero, A. La Greca, S. Bhansali, N. Bourguignon, C. Luzzani, M.S. Pérez, S. Miriuka, B. Lerner

## Abstract

Microfluidic tools have recently made possible many advances in biological and biomedical research. Research fields such as Physics, Engineering, Chemistry and Biology have combined to produce innovation in Microfluidics which has positively impacted on areas as diverse as nucleotide sequence, functional genomics, single-cell studies, single molecules assays, and biomedical diagnostics. Among these areas regenerative medicine and stem cells have benefited from Microfluidics due to these tools have had a profound impact on their applications. In the study, we present a high-performance droplet-based system for transfecting individual human-induced pluripotent stem cells. We show that this system has great efficiency in single cells and captured droplets, similar to other microfluidic methods and lower cost. We demonstrate that this microfluidic approach can be associated with the PiggyBac transposase-based system to increase its transfection efficiency. Our results provide a starting point for subsequent applications in more complex transfection systems, single-cell differentiation interactions, cell subpopulations, cell therapy, among other potential applications.

## 1. Introduction

The field of regenerative medicine has been radically moved forward with the development of human induced pluripotent stem cells (hiPSCs). These cells are obtained by genetically reprogramming terminally differentiated adult cells, and can be later maintained indefinitely in *in vitro* culture and can be differentiated to all the cell types in the adult organism^1^. Thus, these cells provide unprecedented opportunities to study the earliest stages of human development *in vitro*, to model various human diseases, to perform drug tests in culture and, ultimately, as an unlimited source of cells for future therapeutic applications. To take advantage of this potential, it is essential to be able to control the differentiation of hiPSCs to somatic lineages with high efficiency and reproducibility in a scalable and economical way^2,3^.

Cell-to-cell heterogeneity has been an important discussion factor in the phenotypic-genotypic profile of healthy and diseased tissues^4,5^. Single cell analysis technologies seek to study this heterogeneity in order to understand cellular, molecular and tissue physiology. In contrast to bulk tests, heterogeneity measurements through individual cell assays provide a much higher phenotypic resolution and do not require prior knowledge of cell subtypes within a sample^6^. Single cell behavior depends on the properties of complex niches that provide a wide variety of biochemical and biophysical signals^7^. In order to unravel and understand the behavior of these cells it is necessary to deepen the analysis even to see the unicellular behavior and the possible cellular subpopulations that exist within apparently homogeneous tissues.

A rapidly developing field that has proven to be very useful in single cell analyses is microfluidics. Microfluidics refers to the science that allows the study and manipulation of fluids in the micrometer scale^8^. Over the last years, microfluidic systems have been established as promising platforms for many lab-on-chip (LOC) applications in many fields of research, such as chemistry, biology, medicine or engineering^9,10^. Their main advantages lie in the ability to control liquids under the laminar regime, reduction in the number of reagents and samples, shorter analysis time, reduction of costs and portability. Polydimethylsiloxane (PDMS) is the most used material for the manufacture of microfluidic microdevices because it is optically transparent, biocompatible and easy to manufacture. Droplet microfluidic systems allow the isolation of individual cells in conjunction with reagents in liquid capsules or picolitre monodisperse hydrogels with a yield of thousands of droplets per second. These qualities allow many of the challenges in the analysis of a single cell to be overcome. Monodispersity allows quantitative control of reagent concentrations, while droplet encapsulation provides an isolated compartment for the single cells and its environment. This high performance allows the processing and analysis of the tens of thousands of cells that must be analyzed to accurately describe a heterogeneous cell population in order to find uncommon cell types or access sufficient biological space to find successes in a directed evolution experiment^11^.

Functional genomic studies such as the transfection of fluorescent reporters are essential for unraveling cellular and molecular biological phenomena^12^. When trying to generate stable cell lines that incorporate a construct within the genome, the PiggyBac technology based on the system of transposable mobile genetic elements has recently gained ground as an alternative to classical methods such as viral transduction. Briefly, this methodology relies on the transient transfection of a plasmid with the sequence to be inserted flanked by inverted terminal repeat sequences (ITRs) and a transposase enzyme that catalyzes the integration of the foreign DNA between AATT chromosomal sequences. This methodology has proven highly efficient in human and other mammalian cell types^13,14^

In response to this, numerous techniques have been generated in order to increase the efficiency of transfection^15^. Individualizing the cells and enclosing them within droplets at the picolitre scale allows the cell to have greater contact with the reagents and plasmids, consequently increasing the transfection efficiency. However, production of hundreds or thousands of droplets can limit the optimal individual follow-up.

In this work, we develop a microfluidic system of two interconnected microdevices that have a high performance at the time of individualizing the cells and a low manufacturing cost compared to similar systems^16,17,18^. The latter often use SU8 resin as the manufacturing base which is significantly more expensive and a single microdevice to form the droplets and store them this tends to limit long-term cultivation and future clonal selections. Unlike the methods described above, our method of manufacturing the storage microdevice uses a PDMS multilevel system, which allows the droplets to be stored according to manufacturing size and depth. The process consists of two stages. First, a microdevice that is responsible for generating monodisperse droplets in which the cells (in our case, hiPSCs) were captured together with the plasmids and transfection reagents. The second part consists of a multilevel microdevice designed with thousands of wells where the droplets are stored allowing cell culture and real-time monitoring of cell-to-cell transfection and cell viability over time. In addition, the intrinsic advantage of being separate microdevices allows long-term cultivation and multiple uses of accurate cell visualization tools. We strongly believe that relatively cheap devices could function as a starting point for subsequent applications in more complex transfection systems, unicellular differentiation interactions, or cellular subpopulations, among other potential applications.

## 2. Materials and methods

### 2.1 Design and fabrication of microfluidic microdevices

Two separates microdevices were designed for droplet formation and storage droplet applications. Each of these were manufactured with different methods in order to improve system versatility. The microdevices were manufactured as described previously^19^, in the case of the forming microdevice of droplets. Briefly, a mold in high relief with the desired design was made by photolithography in a 700 μm thick silicon wafer (Virginia Semiconductor, Inc.), using the negative resin SU-8 (MicroChem). The microchannels have a final height of 150 μm. Next, the mold was placed under vacuum with trichloro (1H, 1H, 2H, 2H-perfluoro-octyl) silane (Sigma) for 1 hour, to protect the SU-8 resin from detachment by releasing PDMS from the mold. The PDMS was mixed with the curing agent in a 10:1 ratio and the mixture was placed under vacuum for 1 hour to remove air bubbles. Next, the mixture was poured back under vacuum for 1 h, and cured in an oven at 70 ° C for 70 min. The PDMS was not molded and the fluidic connection ports were constructed by drilling holes in the PDMS with a syringe needle (21-gauge, internal diameter of 0.51 mm). Finally, the PDMS was assembled with the glass base. Through the plasma oxygen system (deposition of chemical vapors enhanced with plasma), the microdevice and the glass base are exposed for 3 minutes at a pressure of more than 4000 g overnight.

In the case of the droplet storage microdevice, a new custom made multilevel microfluidic manufacture technology was used^20^. Briefly, the droplet storage microdevice was designed with a Layout Editor design editor software and transferred to the TIL with a 2400 ppi infrared laser source, then the TIL was laminated on the unexposed photopolymer plate, the photopolymer plate was exposed at UVA light at 0.45 J on the back for 10 s, a part of the photopolymer was covered with a mask plate on the back side, the photopolymer plate was exposed to UVA light at 0.45 J on the back side for 20 s, the previous exposures were repeated one at a time, finally the front side was exposed to UVA light at 19 J for 360 s, after the TIL was removed, the plate was washed with PROSOL N-1 solvent (supplied by Eastman Kodak) at 360 mm min −1 was dried in an oven for 30 minutes at 50 °C, the plate had a last exposure to UVC light at 10 J for 17 minutes and UVA light was applied at 4 J for 2 minutes on the front side With the mold manufactured briefly, a mixture of epoxy resin and curing agent (Cristal-Tack, Novarchem – Argentina) was poured over the female mold to replicate the design in high relief.

After curing, the epoxy resin mold (ERmold) was detached from the Fmold to form the male mold. Subsequently, a mixture of PDMS and curing agent in a 10:1 weight ratio (Sylgard 184 silicone elastomer kit, Dow Corning) was poured onto the ER mold and cured in an oven at 40 °C overnight. The same steps described for the droplet forming microdevice were used for assembly.

### 2.2 Production of droplets storage and breaking

For the production of monodisperse droplets, the biocompatible oil FluoroSurfactant-HFE 7500 5% was used for the continuous phase, and for the dispersed phase the cells were used in suspension in mTeSR™l (STEMCELL Technologies) medium with Rock Inhibitor (Y-27632). The injection of the phases was performed using the infusion set Fullgen A22 (USA), with results for the continuous phase of 4.50μl / min – 6.50μl / min, and for the dispersed phase 1.75 μl / min – 3.00μl / min. The average size of the droplets generated was 80 μm – 87 μm, then a tube was connected to the output hole of the producer microdevice and the inlet hole of the droplets storage microdevice was progressively increased to complete the storage volume.

In order to release the cells that were encapsulated within the oil droplets. SIGMA 1H, 1H, 2H, 2H perfluoro-octanol (PFO) was used based on the protocols recommended^21^.

### 2.3 Droplet images

Nikon SMZ645 Stereo Zoom Microscope and a Canon T3-I Rebel digital camera connected to the microscope were used to capture the droplets. The images were created from a stack of multiple acquisitions of microscopes in a large surface area of the microdevice.

The analysis and quantification of the results obtained in the experimental phases were performed using the ImageJ software and R.

### 2.4 2D Cell culture

The hiPSC cell line used in this study was generated in our lab from adult male fibroblasts and was previously described^22^. Cells were regularly cultured in 6 well plates in E8-Flex medium with Geltrex coating (all Thermo Fischer Scientific). Cells were passaged every 2-3 days using Tryple until the moment of encapsulation and subsequent transfection. All of cells cultures were maintained at 37 °C in a saturated atmosphere of 95% air and 5% CO2.

### 2.5 Plasmids used in this study

To assess the transfection efficiency and to generate stable cell lines two plasmids were used^23^. A first plasmid that encodes a CAG-mCerulean-H2B reporter together with a Neomycin resistance cassette, both flanked by ITR sequences. The second plasmid drives the transient expression of the PiggyBac transposase.

### 2.6 Cell viability and nuclear staining

To measure cell viability after drop encapsulation and transfection, we used the Live Dead Assay kit (Thermo). Since the drop-based system differs according to standard culture techniques, we empirically determined that 5 times more reagent was required than recommended by the manufacturer. On the other hand, Hoechst 33342 was used for staining cell nuclei in living cells in order to measure transfection efficiency.

## 3. Results and discussion

### 3.1 Microfluidic system design

The constructed single cell microfluidic transfection system is illustrated in Figure 1A. This system consists of two interconnected microdevices and an outlet channel. The former produces monodisperse droplets containing cells, plasmids, and transfection agents using the flow-focusing system. These flows are controlled by infusion pumps with infusion and pressure control. The second microdevice is made up of two layers of PDMS that have 3000 multi-level wells that allow a high droplet capture performance and an output channel to recover them. In addition, this double-layer design allows for greater gas exchange in conjunction with fluorinated oil (HFE-7500), which greatly helps cell viability^24^.

**Figure 1.**
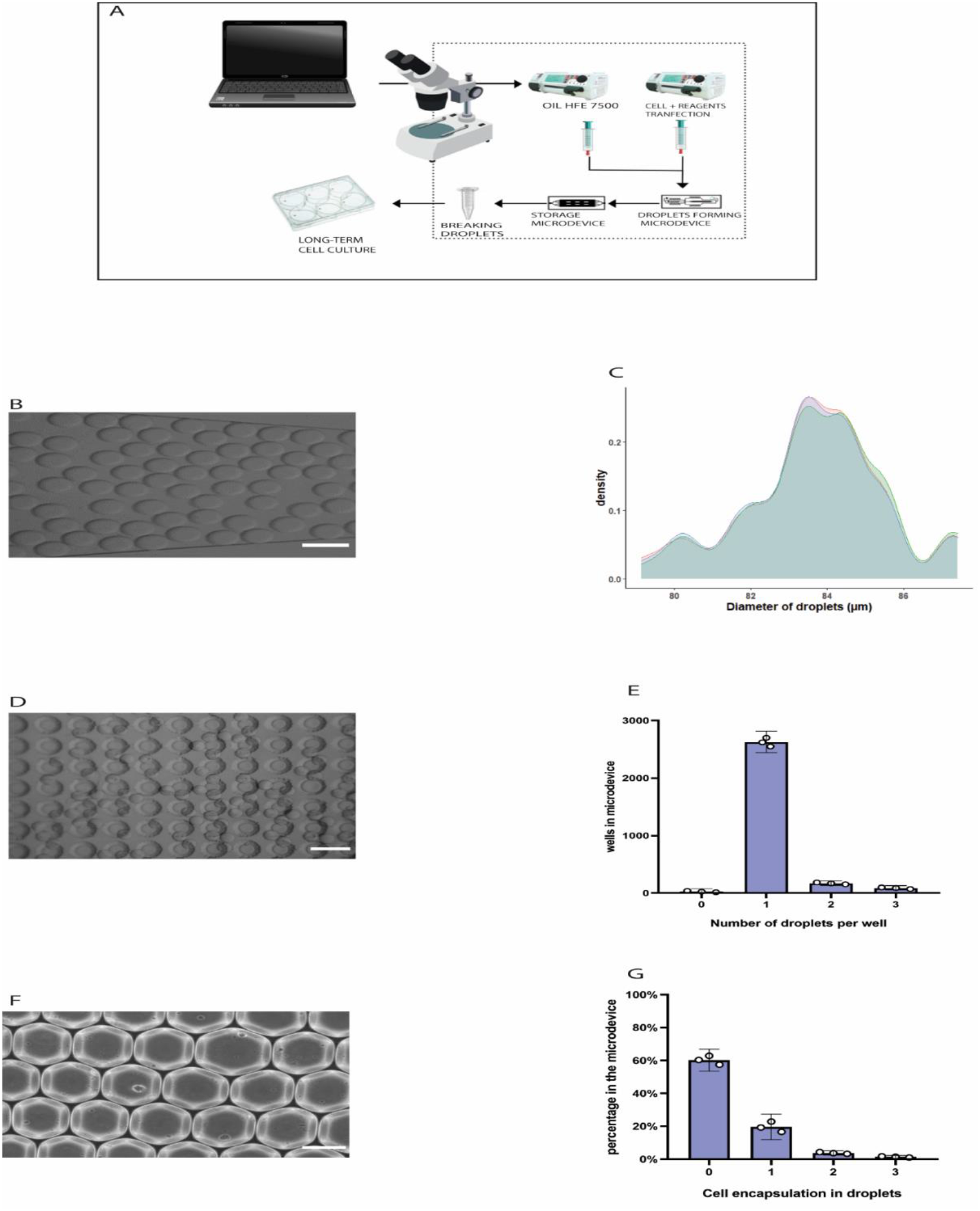
Characterization of the microfluidic system. A. The scheme shows a diagram of the single cell microfluidic transfection system for hiPSC. B. Representative image of the droplets produced in the outlet channel of the forming microdevice. C. The density graph shows the size of the droplets produced by the system. Each line represents each technical sample that was made. D. Representative image of droplets captured in the micro storage device. E Number of droplets captured per well of the micro storage device. White points indicate the mean value for each replicate (N = 3). F. Representative image of cells inside that travel through the outlet channel of the forming device. G. The bar graph shows the effectiveness of capturing a cell per droplet. White points indicate the mean value for each replicate (N = 3). The scale bars correspond to 100 μm.

### 3.2 Droplet production, storage and encapsulation

First, the efficiency of the microdevices was characterized that conform our system. Figure 1B shows the production of droplets in the outlet channel of the forming microdevice. A total of 1000 droplets average size of 83 μm (Figure 1C). The perfusion rate was 6.50 ul / min for the continuous phase (oil) and 3.00 ul / min for the dispersed phase (cells, plasmids and transfection agents).

On the other hand, the efficacy of the microdevice that stores the droplets was measured (Figure 1E). High efficiency is determined by the storage microdevice to capture one droplet per well. This helps a lot in image analysis and real-time cell detection.

In Figure 1D the wells of the storage microdevice are observed capturing the droplets with cells inside.

We next evaluated the level of encapsulation of cells per droplet. Figure 1F shows cells encapsulated within monodisperse droplets formed in the system. As expected, most of the droplets did not contain any cells, while we observed and efficiency of 22% in single cell encapsulation (Figure 1G). These values resemble similar work of encapsulation and storage of droplets^25^.

### 3.3 Cell viability in the microfluidic system

It is widely known that hiPSC cells are highly sensitive to changes in their environment that can reduce their viability and event prompt them to enter pathways of apoptosis^26^.First, in order to demonstrate the effectiveness of our system viability cells were evaluated at different times after encapsulation and subsequent release of droplets. For this purpose, used the methodology of the Live Dead test kit, which emits different fluorescent signals depending on the state of viability of the cells. (Figure 2A). We observed that there was a high cell viability at 8 and 16 hours. In contrast, the longer the cells encapsulated in the droplets spent, their viability decreased significantly. At 24 hours of encapsulation and at 48 hours, a high degree of cell death and low viability with time was evident (Figure 2B).

**Figure 2.**
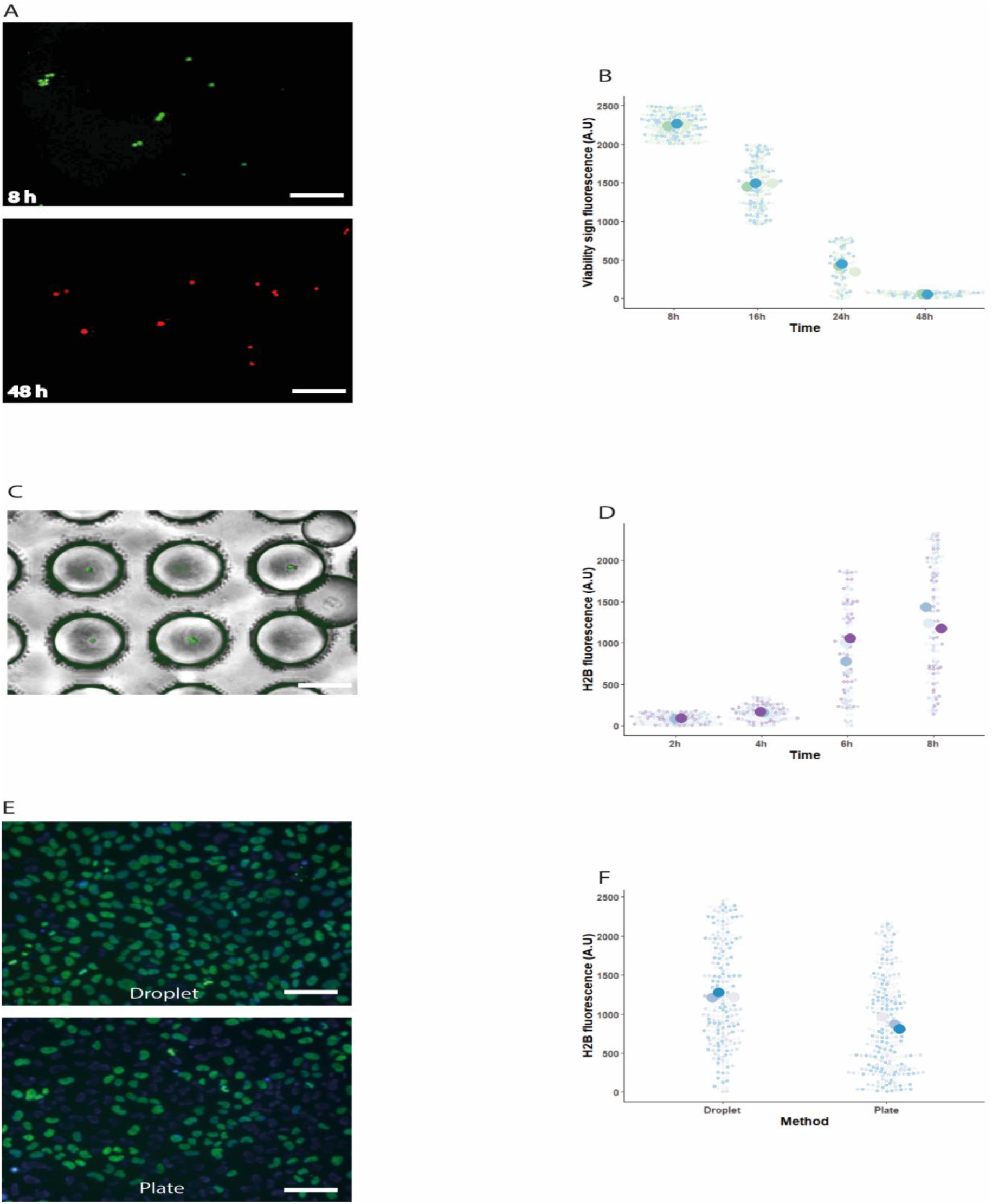
Transfection and method efficiency. A. Representative image of cell viability. Green fluorescence indicates cells with marked cell viability, while red fluorescence shows cells that are not viable or in an apoptotic state B. Cell viability as a function of encapsulation time. Small points correspond to mean intensity measurement of individual cells within each biological replicate as indicated by different colors (N=3). Larger points depict mean value for all measurements in each specific biological replicate. C. Representative image of cells within the droplets. Each drop is captured in a well of the storage microdevice. D. H2B-mCerulean fluorescence at different times after encapsulation with transfection reagents. E. The images show the comparative top platelet of the cells transfected into droplets and subsequently released. The image below corresponds to the traditional culture plate transfection method. F. Comparison of H2B-mCerulean transfection in the microfluidic system or in multiwell plates. In the microfluidic system, cells were encapsulated with transfection reagents and the reporter plasmid for 8h, droplets were broken and cells were later cultured in multiwell plates until fluorescence was measured 24h after encapsulation. In the case of multiwell transfection, reagents were incubated during 8h, medium was changed and fluorescence was evaluated at 24 h. The scale bars correspond to 200 μm.

### 3.4 Efficient transfection of hiPSCs using the microfluidic system

Cell transfection is an invaluable tool as it allows the generation of engineered cell lines with Crispr/Cas9, the overexpression of proteins, gene silencing, fluorescent reporter visualization, among many other. Due to hiPSCs display high cell viability during the first 8 hours after encapsulation, we decided to evaluate the transfection efficiency of the cells within the droplets. Therefore, transfection of a fluorescent H2B-mCerulean indicator was evaluated when the transfection reagents were provided during the encapsulation step.

After cell encapsulation and droplet storage, the microdevice was placed under culture conditions at 37 °C in an atmosphere saturated with 95% air and 5% CO2. To assess transfection efficiency, the amount of fluorescence emitted by each individual cell within the droplet was measured in a period of time of 2, 4, 6, 8 hours (Figure 2C). As expected, fluorescence intensity increased with time after encapsulation, with higher levels at 6 and 8 hours. This observation could be explained both because of the time it takes for fluorescent proteins to be transcribed, translated and matured, as well as by the increased likelihood of cell transfection when cells are confined within the droplets with the transfection reagents. Indeed, our microfluidic system is suitable for single cell transfection and that it could be potentially applied in different approaches.

### 3.5 Comparison of plate methods with the microfluidic system

To further characterize our microfluidic transfection setup, the system was compared with conventional plate transfection method in multiwell plates. As shown in the previous transfection experiment, the optimal encapsulation time taking into account cell viability and transfection was 8 hours. In this context, we compared both methodologies with the same transfection time and with the same initial concentrations of plasmids and lipofectamine.

Figure 2E shows the comparison of both methods. As there is a significant proportion of transfected cells in both strategies, results indicate a significantly higher intensity level of fluorescence in the microfluidic system (Figure 2E-F), which could be a consequence of the confinement of the transfection reagents within the droplets.

Although cell viability was reduced after prolonged culture within the droplets, our results show that transfection of hiPSCs in our microfluidic system was comparable and even displayed higher levels of transfection compared to the conventional methodology in multiwell plates.

## 4. Conclusions

The developed droplet-based microfluidic system showed high transfection efficiency of HiPSC cells, by inserting the H2b-cerulean fluorescent protein. We hypothesized that this could be due to the microenvironment conditions cells were exposed to at encapsulation time along with the plasmids and the transfection reagents.

The microfluidic system also proved to be highly effective in capturing one droplet per well and individualizing cells within the droplets for subsequent real-time monitoring. This is attributed to the multi-level storage microdevice, which allowed visualization and location of each cell during the transfection process.

Single cell transfection performed by microfluidic droplet method was shown to be highly effective and economical compared to other similar methods. We believe that our microfluidic system could be a powerful tool for assays where isolation and transfection of individual cells are needed, such as the generation of clonal cell lines using Crispr / Cas9 technology, regenerative medicine and gene therapy by providing homogeneity by transfecting individual cells. Beyond the current study focused on gene editing by transfection, this innovative system can be used for various purposes. Such as the production of proteins from individual cells, microbioreactors, selective isolates, drug testing, among other applications at costs that most laboratories could access.

## Acknowledgements

The authors thank the financial support from CONICET (IP2015), ANPCyT (PICT-STARTUP 4819) and Florencio Fiorini grant. We would like to thank Jorge. L. Fernandez and I. de Sá Carneiro for general support and fruitful discussions.

## Notes

### Competing Interest Statement

The authors have declared no competing interest.

